# Direct Visualization of Anesthesia-induced Subcellular Dysfunction in Non-Neuronal Tissues of C. elegans

**DOI:** 10.1101/2025.07.03.663001

**Authors:** Bodhidipra Mukherjee, Shilpa Chandra, Abdul Salam, Laxmidhar Behera, Chayan Kanti Nandi

## Abstract

Volatile anaesthetics such as isoflurane, halothane, and sevoflurane are extensively utilized for their reversible induction of unconsciousness, exhibiting well-defined impacts on neural networks. However, their influence on non-neuronal and developing tissues is inadequately comprehended, generating concerns for vulnerable categories, including children and individuals receiving extended anesthesia. To rectify this deficiency, we examined the subcellular impacts of anaesthetics utilizing HEK 293T and HepG2 cell lines, alongside with Caenorhabditis elegans as a comprehensive organismal model. In vitro treatment to lidocaine and isoflurane resulted in mitochondrial depolarization, lysosomal aggregation, and buildup of reactive oxygen species (ROS). In vivo, 8% isoflurane induced mitochondrial fragmentation, loss of branching, and disruption of tubular lysosomes, indicating compromised energy metabolism and autophagic stress. We detected selective transcriptional inhibition of neuron- and immune-related promoters, although RNA Pol I remained active, indicating energy-conservation strategies. Systemic oxidative stress was validated by TIR-1::GFP aggregation and lipofuscin accumulation. Our research is one of the initial investigations that defines anaesthesia-induced subcellular dysfunction beyond the nervous system, providing novel insights for safer anaesthetic protocols and informing long-term health evaluations, especially in individuals who are developing or chronically exposed.

**Significance statement:** While volatile anaesthetics are widely used for inducing unconsciousness, their effects beyond neurons remain poorly understood. This study reveals that clinically used anaesthetics such as isoflurane and lidocaine cause profound subcellular dysfunction in non-neuronal systems, including mitochondrial depolarization, lysosomal disruption, transcriptional repression, and systemic oxidative stress. Using both human cell lines and *Caenorhabditis elegans*, we uncover conserved patterns of organellar damage and stress responses. These findings have critical implications for pediatric, intensive care, and long-term anaesthesia cases, urging the need to reassess anaesthetic safety profiles and develop biomarkers for systemic toxicity. Our work opens new avenues for designing cell-type-specific protective strategies under anaesthetic exposure.

## Introduction

Volatile anaesthetics like isoflurane, halothane, and sevoflurane have long been used in medical practice because of their efficacy in inducing a state of reversible unconsciousness. This trait becomes widely effective during procedures requiring pain elimination or reducing inflammation after accident recovery or surgery [1]. Since their first effective usage in the 1840s by William T.G. Morton who used diethyl ether at Massachusetts General Hospital, it led to the age of volatile anaesthesia and the beginning of their gradual adoption in more complex and longer duration surgeries [2]. However, researchers soon discovered that diethyl ether was highly toxic, leading to nausea, vomiting, and more severe side effects after long-term exposure. Hence, throughout the 19th and 20th centuries, research was conducted on developing a class of these volatile anaesthetics that show lower toxicity and more controllable kinetics [3]. Volatile anaesthetics have several advantages, for example, they have low blood and tissue solubility, leading to more rapid onset and recovery from their effects, and are also eliminated from the body predominantly via exhalation, allowing for more rapid control of their kinetics [4]. Their method of delivery is non-invasive via ventilation, reducing patient stress, and because they are eliminated from the body mainly by exhalation, they show lower hepatotoxicity [5, 6].

The mechanism of action of volatile anaesthetics focuses chiefly on their effect in neuronal cells, where they inhibit voltage-gated sodium channels, which prevent the propagation of action potentials [7], reduce calcium influx via NMDA receptors in the glutamatergic neurons, which prevents neurotransmitter release and propagation of nerve impulses in the primary excitatory pathway [8] and also increase the duration and frequency of opening of GABAA receptor channels, which causes chlorine influx and hyperpolarizes the neuron, making it less likely to fire action potentials [9]. However, the effect of repeated anaesthetic exposure or anaesthetic overdose on more susceptible individuals, like children, is not properly understood. Volatile anaesthetics can damage mitochondria in stressed or developing cells, they primarily inhibit complex 1 of the mitochondrial electron transport chain (ETC), which leads to disruption of the ETC and reduced ATP production [10]. A disrupted ETC can cause electron leakage, which in turn generates ROS that damages mitochondrial DNA, proteins, and lipids and activates autophagy [11]. Such cellular stresses can trigger the release of caspases and the apoptosis [12] of newly formed or developing neurons, which is especially harmful in children.

Most of the studies related to anaesthetic effect focus primarily on neurons with no detailed study existing that reports their effect on other cells in the body. In a recent study, the authors showed that exposure to sevoflurane caused serious harm to glial cells, especially oligodendrocytes, in newborn mice. However, there is little emphasis on how these anaesthetics may affect other non-neuronal cells. Given the lipophilicity of volatile anaesthetics like isoflurane or halothane [15], they have the ability to cross the lipid membrane of a cell and get absorbed anywhere within the body. The human body is a large entity made up of a complex mass of different types of cells, and a single exposure to an anaesthetic effect may not be able to cause detrimental changes in the body. However, for patients who are repeatedly exposed to anaesthesia or in more susceptible individuals like children, we do not know the changes an exposure or an overdose may cause. Obtaining real-time data regarding this phenomenon from a human being is not possible. Hence, in this study we investigate a combined in vitro and in vivo approach. The HEK 293T and HepG2 cell lines were employed to evaluate initial organellar responses to lidocaine and isoflurane, whereas Caenorhabditis elegans functioned as a whole-organism model to investigate systemic and chronic effects. In vitro, anaesthetic treatment resulted in mitochondrial depolarization, lysosomal aggregation, and reactive oxygen species buildup, signifying early stress. In C. elegans, prolonged exposure to 8% isoflurane (v/v) resulted in mitochondrial fragmentation, lysosomal degradation, activation of the stress gene TIR-1, and downregulation of immunological and thermosensory genes [16,17]. Nuclear morphology underwent significant alteration, becoming significantly round, presumably due to cytoskeletal injury. Collectively, our findings demonstrate that anesthetics that are volatile cause extensive subcellular disruption, including mitochondrial function, autophagy, transcription, and nuclear integrity. By clarifying these pathways, our study enhances the comprehension of how anaesthesia influences non-neuronal systems, informing our development of safer anaesthetic systems, particularly for pediatric and neurocritical care. These findings further support for the development of individualized anaesthesia protocols for individuals necessitating recurrent exposure, so enhancing patient safety and mitigating long-term adverse effects.

## Results

For our work, different strains of C. elegans were used, with GFP tags on specific organelles to show changes in their behavior over time. C. elegans were initially grown in nematode growth media (NGM) and fed E. coli OP50 strain **(Figure S1)**. The animals were age-synchronized via bleaching (discussed in the methods section); L2-L3 stage animals were chosen for experimentation. Several strains of C. elegans were chosen for The animals were exposed to 8% isoflurane, and time-dependent changes were observed. We used a variety of C. elegans strains that had fluorophores tagged to specific organellar proteins for the experiments. To understand mitochondrial dynamics, we used C. elegans strain GL390, which has a C-terminal GFP tag inserted into the endogenous aco-2 locus [18]. Aco-2 is a mitochondrial marker, and the tag allows for visualization of the organellar dynamics upon treatment. For visualization of lysosomal dynamics, we used strain XW8056 which expresses scav-3::GFP [19]. Scav is a lysosomal membrane protein and the tag allows for long term visualisation of lysosomal dynamics. Strain ESC360 with GFP tag inserted at the N-terminus of the endogenous rpoa-2 locus and constitutive expression of myo-3p::TIR1::mRuby were used for visualisation cellular stress in muscles and monitor RNA polymerase activity [20]. Strain AU78 was used for visualisation of transcriptional efficiency of TTX-3 promoter and T24b8.5 promoter under anesthesised condition [21].

### 1. Effect of Anaesthesia In-Vitro

Mitochondria are some of the most important organelles in the cells of an animal’s body. Not only do they carry out most of the oxygen-based energy production in the cell through the Krebs cycle and electron transport chain (ETC) [22], but they also help reduce stress in the cell by controlling calcium levels and producing heat to keep body temperatures stable [23], [24]. Anaesthetics according to previous literature have been shown to cause organellar dysfunction, one of the primary organelles targeted by anaesthesia are mitochondria where the molecules directly block Complex 1 of electron transport chain (ETC) [25]. This may lead to electron leakage generate ROS and cause more mitochondrial damage in a positive feedback loop. To test mitochondrial viability in-vitro under anaesthetic conditions we used HEK 293T cells and treated them using liquid anaesthetic Lidocaine at 0.02% solution concentration. Mitotracker green, (MTG) was used to label mitochondria and observe their dynamics under anaesthesised state. We observed a huge decrease in mitochondria related fluorescence overtime in the HEK cells when they were exposed to the anaesthetic, (**Figure 1 a, I-V**) something not observed in control cells (**Figure 1 b, I-V**). The results justify large scale dysfunction in mitochondrial viability within the HEK 293T cells after anaesthetic application. ROS due to electron leakage from mitochondria can also effect lysosomal behaviour since ROS may trigger protein misfolding cause organellar damage by lipid peroxidation [26] it may trigger autophagy and cause changes in number and positioning of lysosomes. In (**Figure 1 c, I-V**) we see that in control the large well formed lysosomes are distributed evenly all over the cells, compared to the lidocaine exposed samples (**Figure 1 d, I-V**) where we observe a sudden decrease in in their number in the beginning of the treatment followed by their time dependant aggregation in certain cellular areas. The results signify that anaesthesia may disrupt proper lysosomal functioning at cellular level.

**Figure 1:**
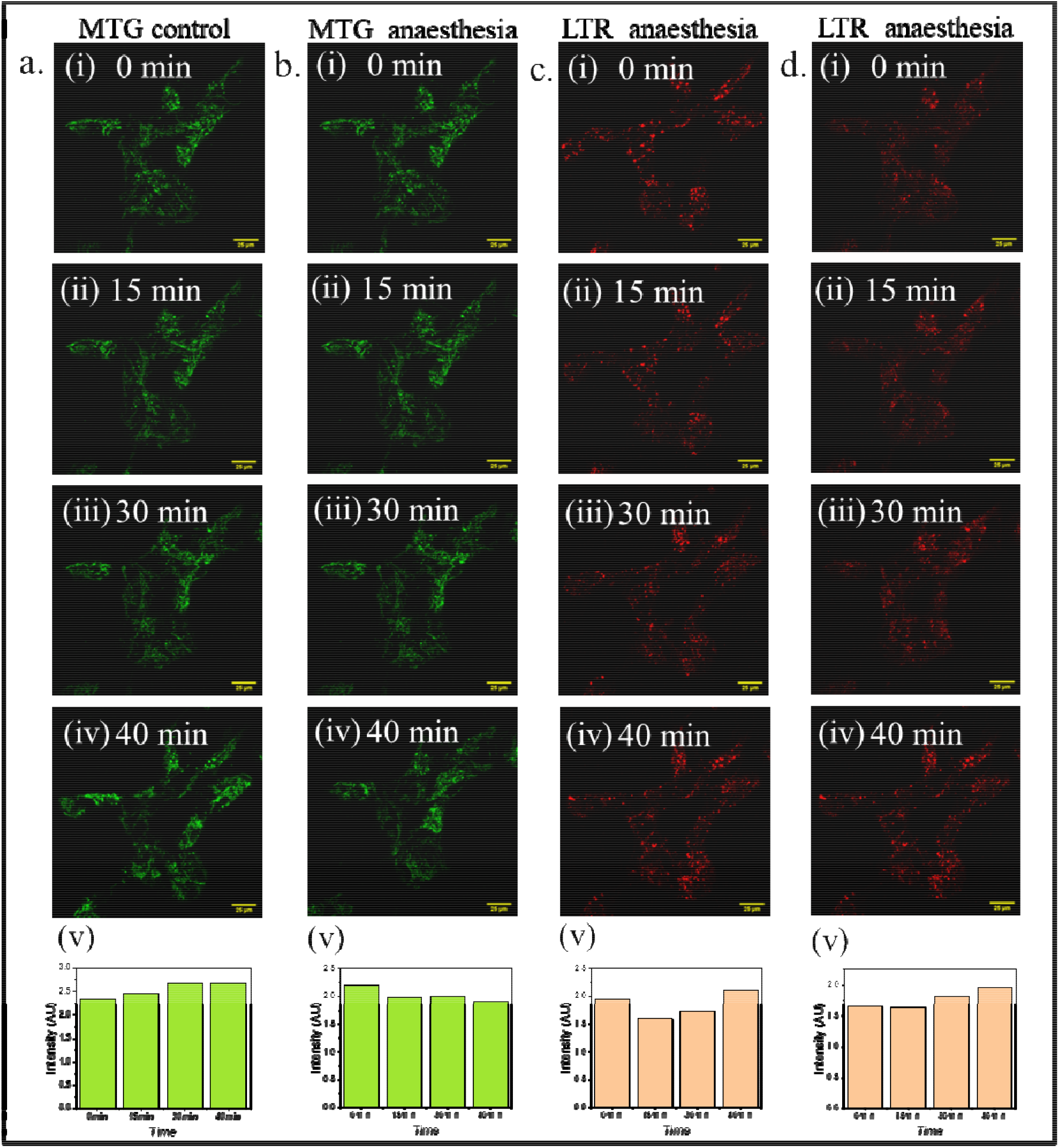
Effect of anaesthesia on HEK293T cells. **a (i-iv):** Not much visible change in MTG based fluorescence in control HEK293T over time. **a (v)**. Overtime intensity change plot. **b (i-iv)**. Under anaesthetised condition HEK 293T cell showed time dependant decrease in fluorescence. **b (v)**. Overtime intensity change plot. **c (i-iv)**. Not much visible change in LTR based fluorescence in control HEK293T over time. **c (v)**. Overtime intensity change plot. **d (i-iv)**. Under anaesthetised condition HEK 293T cell showed time dependant increase in LTR fluorescence. **d (v)**. Overtime LTR intensity change plot

#### 2. Effect of Anaesthesia In-vivo

#### Effect of anaesthesia on mitochondria

The invitro experiments revealed that anaesthetic exposure may effect non neuronal cell lines which led us to study its effects in live animal systems like C. elegans. Given the vast number of important works done by mitochondria to maintain optimal organismal health, we decided to check its health and any changes associated with it upon inducing a state of unconsciousness with anaesthetics. We used C. elegans with GFP tagged to mitochondrial proteins for the observations. We found a temporal change in mitochondrial structure throughout the entire body of a C. elegans when they were exposed to 8% isoflurane, as seen in **(Figure 2 A-C)**. The average area and perimeter of mitochondria decreased **(Figure 2 D,E)**. This observation can be correlated to mitochondrial fragmentation due to cellular dysfunction [27], pointing to the fact that anaesthesia can trigger high levels of stress on the applicant after administration. Anaesthetic treatment also triggered increased circularization of the mitochondrial structures **(Figure 2F)**. In a healthy state, mitochondria exist as long tubular structures, but under high stress conditions, dysfunctional mitochondria can assume a circular shape, further strengthening the idea of anaesthesia induced mitochondrial dysfunction. Healthy mitochondria look like a network of connected tube-like structures with many junctions [28]. We also observed from our experiments a time-dependent reduction in the number of junctions after application of anaesthesia **(Figure S2)**, implying a global collapse in their viability and ability to function effectively.

**Figure 2:**
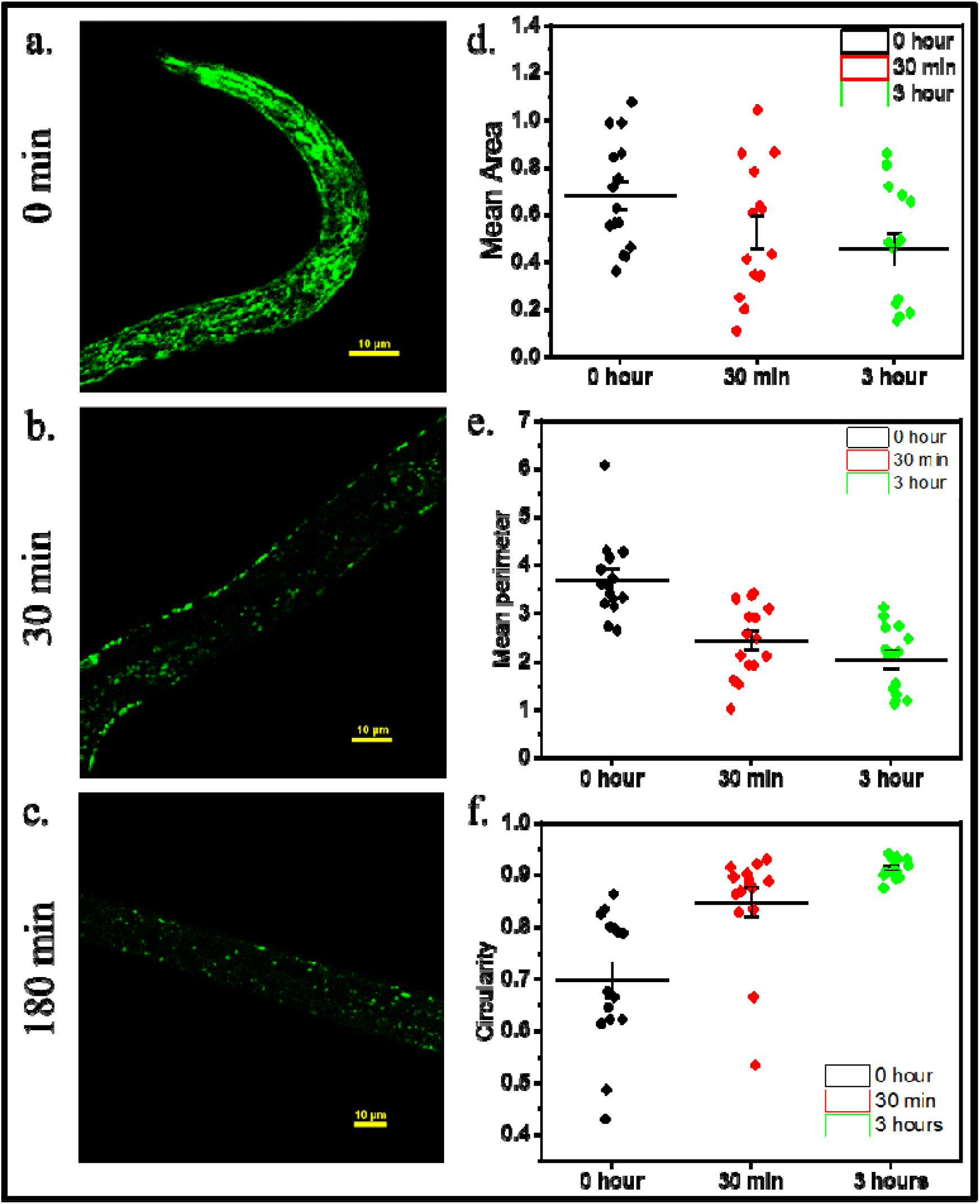
Effect on anaesthetics on C. elegans mitochondria. **(a-c)**. Temporal decrease in mitochondria related fluorescence observed in GL 390 strain of C. elegans after treatment with an aesthesia.(**d)**. Decrease in overall mitochondrial area over time after isoflurane treatment. **(e)**. Decrease in perimeter of all mitochondria in C. elegans after being anaesthetised with isoflurane. **(f)**. Increase in overall circularity of mitochondria in C. elegans after being subjected to anaesthetic treatment.

#### Effect of anaesthesia on lysosomes

Lysosomes are an important organelle that maintains optimal organismal health and performs multiple functions, from uptake of damaged dysfunctional organelles by autophagy [29] and aiding in cleanup to storage of metals like zinc in lysosome-related organelles (LROs) [30].Hence, we decided to check lysosomal health after anaesthesia application. In healthy condition,C. elegans intestinal and epithelial cells can have long tubular lysosomes (TLs) up upto 15μm long as seen in (**Figure 3 A)**. These TLs perform multiple functions, like degradative cargo turnover when initial steps of autophagic machinery are compromised in other lysosomes, expanding the reach of lysosomes to multiple areas of the cell for efficient cleanup of damaged dysfunctional proteins, and have also been associated with enhancing the lifespan of the animal. Our work shows that anaesthetic treastment can reduce the abundance of these TLs in a temporal fashion, as can be seen from the images in (**Figure 3 B-F)** and the data in their subsequent analysis in (**Figure 3 G-I)**. From **(Figure 2, part G)**, we can see that the average area of the lysosomes has decreased from an average of 2.59 μm^2^ in the control to 1.59 μm^2^ after 2 hours of exposure, while in (**part H)** we can see the average perimeter has decreased from 7.1 μm in the control to 5.3 μm after 2 hours of exposure, signifying an overall decrease in size. Over the same time period as seen in (**part I)**, the circularity of the structures has also increased from 0.68 in control to 0.71 after exposure, signifying an overall departure from the tubular morphology of the lysosomes to more circular shapes.

**Figure 3:**
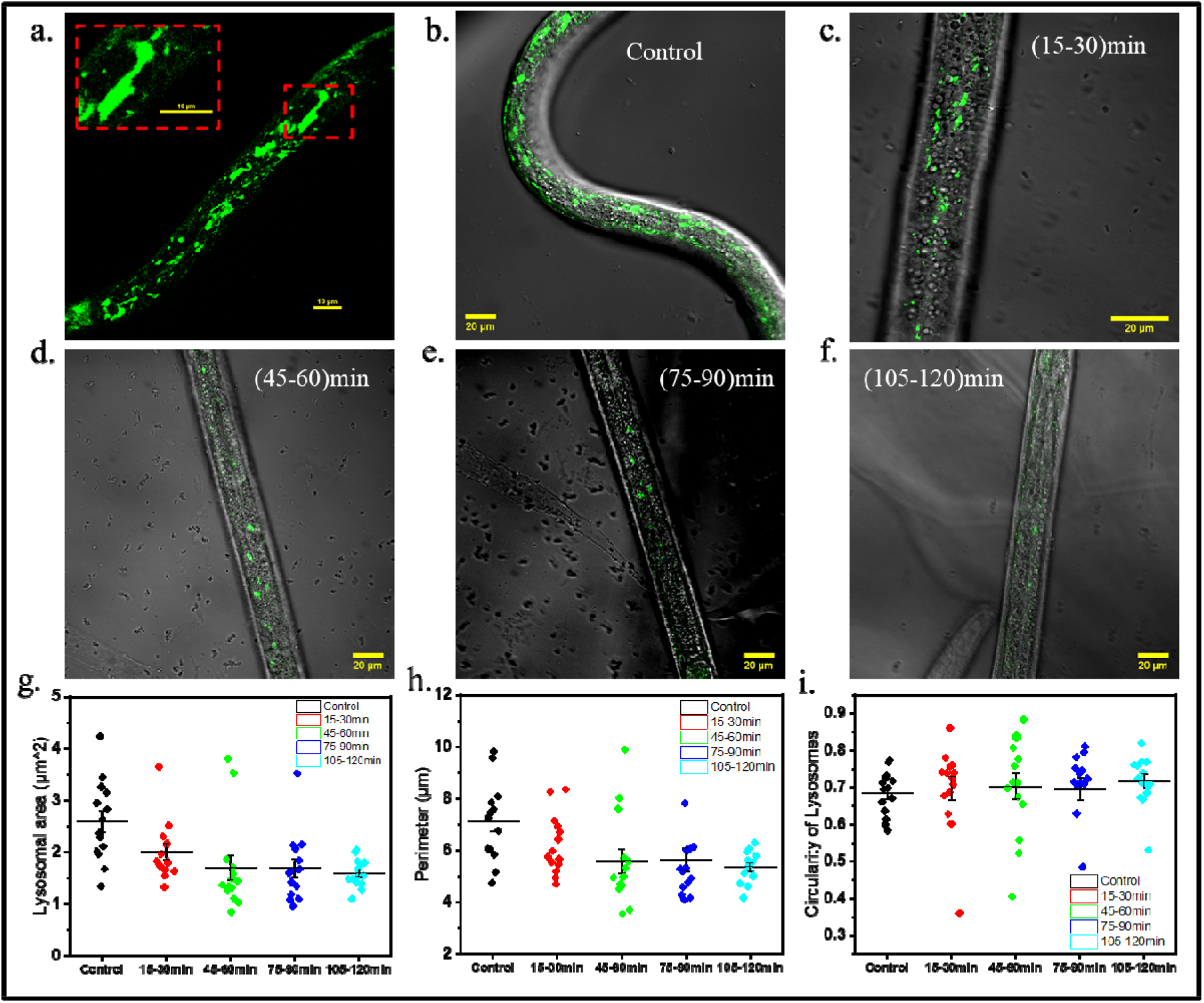
Lysosomal response to anaesthesia induced cellular disruption. **(a.)** Long tubular lysosomes in epithelial region of C. elegans magnified version in top left corner. **(b-f)**. temporal change in lysosomal structure and distribution all over the C. elegans body after being subjected to anaesthetic treatment. **(g-h)**. Time dependant over all decrease in lysosomal area and perimeter observed in C. elegans after exposure to isoflurane **i**. In crease in circularity of lysosomes in C. elegans after they were exposed to anaesthesia

#### Transcription control of anaesthesia

The transcription rates of genes in animal cells are under strict control, some of these controlling elements are called transcription factors. Their function is to aid in upregulation or downregulation of genes and also aid in cell-specific expression of a gene in response to specific external signals or stressors [31]. The ESC 360 strain mentioned earlier has a GFP tag attached to the rpoa subunit of RNA Pol 1, which is a key transcription factor for ribosomal RNA [32]. We observed that the fluorescence intensity associated with RNA pol1 changed sparingly over a 90-hour period **(Figure 4 A-B and C)**, possibly owing to its importance in preserving cellular vitality. Neurons are an essential component of the body of an animal, and in C. elegans, the AIY interneuron is an essential hub for the integration of several sensory inputs like chemosensation, mechanosensation, and behavior modulation in the animal [33]. GFP under the control of the TTX-3 promoter from the AU78 strain was used to check the transcriptional activity levels of this critical neuron cluster. We observed that within half an hour of anaesthetic induction with 8% isoflurane, there was no GFP signal corresponding to reduction in transcriptional activity in these areas **(Figure 4 D-E)**. The same C. elegans strain also expressed intestine-specific GFP under the control of the T24B8.5 promoter. The fluorescence signal from these areas also significantly decreased within half an hour of the same treatment process **(Figure 4 F-G)**. In the body of C. elegans under control of the T24b8.5 promoter, the animal produces proteins bearing homology to ShK toxin peptides [34]. These proteins control immune signaling in relation to pathogen attack [34]. Reduction of expression of GFP from this promoter region may point to a compromised immune response of C. elegans under anaesthesized condition.

**Figure 4:**
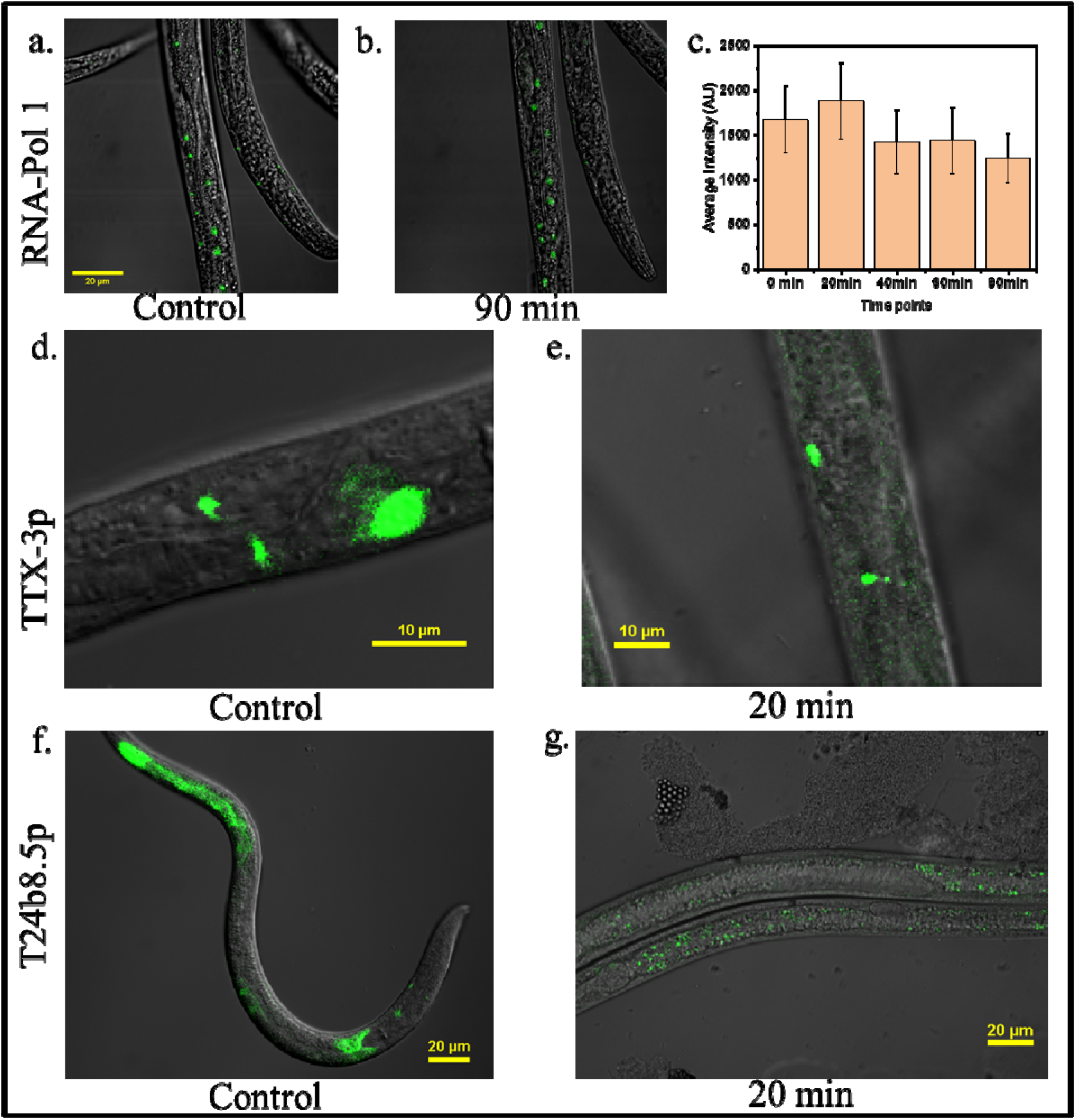
Cellular transcriptional response to anaesthetic treatment in C. elegans. **(a-b)**. Meagre change in RNA-pol 1 based GFP fluorescence observed on ESC360 strain of C. elegans after long term isoflurane exposure. **(c)**. Intensity plot of change in RNA pol 1 based fluorescence showing not much over all change **(d-e)**. Time dependant decrease in GFP fluorescence in AIY neuron cluster from TTX-3 promoter after anaesthetic exposure **(f-g)**. Overall decrease in intestinal fluorescence from T24b8.5 promoter after exposure of C. elegans to isoflurane.

#### Histone dynamics due to anaesthetic exposure

Histones are proteins that provides critical structural support to chromosomes aiding in DNA packaging and regulation of gene expression within a nucleus [35]. In the previous section we observed transcriptional repression of certain immune related and chemosensory genes, since histones play a major role in geneexpression, we decided to see how anaesthetic exposure may affect their biology. The C. elegans GFP tagged with histone 71 was used mainly for the observations and one primary observation we made was a change in the shape of the nucleus of intestinal cells after the C. elegans was exposed to isoflurane. As seen in **figure 5 (a,b)** and their subsequent analysis in **Figure 5 c**, which shows changes in shape in the anaesthesised specimens. The shape being more skewed and irregular in the control animals compared to them being more uniform and circular in shape in animals exposed to anaesthesia. The overall size has also increased considerably. There was also a major decrease in area of histone related GFP fluorescence in the intestinal cells of anaesthesised animals, and most of the decrease can be attributed to the reduction in GFP signal from the nuclei of intestinal cells. As seen in **Figure 5 d**. However interestingly there was no observed decrease in GFP histone related fluorescence in the muscle nuclei. Even after almost an hour of exposure as seen in **figure 5 (e-g)**.

**Figure 5:**
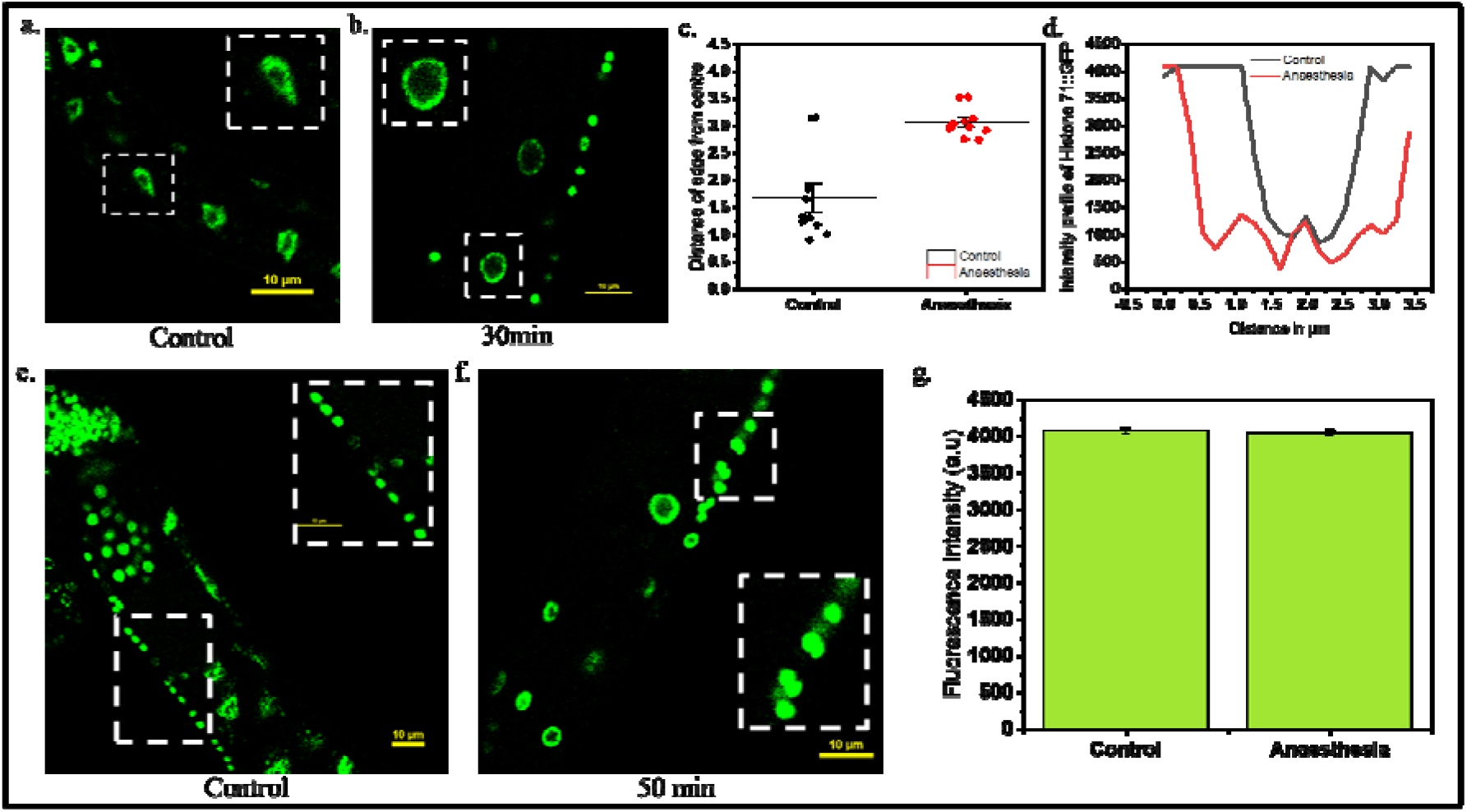
Cellular transcriptional response to anaesthetic treatment in C. elegans. **(a-b)**. Changes in shape of nuclear shape of intestinal cells in C. elegans after exposure to anaesthesia **(c)**. distance of nuclear edge from the centre in intestinal cell **(d)**. Intensity profile of histone 71::GFP related fluorescence in intestinal cells for control and anaesthesised animals **(e-f)**. Intensity of muscle cell nuclei in control and anaesthesised specimens **(g)**. Changes in intensity of histone 71::GFP related fluorescence in muscle cells for control and anaesthesised animals

#### Systemic stress due to anaesthesia

Anaesthesia has been known to cause cellular stress in an animal’s neuronal, endocrine, and immune systems mostly because they induce ROS formation [36]. The generated ROS can trigger protein misfolding, cause ER stress, damage microtubules, and lead to overall cellular dysfunction. In our experiments with 8% isoflurane, ROS levels increased within the entire body of the animal over time, as seen in **Figure 6 (A-E and F)**. The strain used for the study is ESC360, which expresses TIR1-tagged RFP protein in all muscle cells. TIR1 is an important protein that behaves differently based on the types of stress. During pathogen attack, they oligomerize as puncta in intestinal cells [37], during epidermal wounding, their expression levels increase to activate the TIR1/PMK1 pathway to aid in wound closing. In our case we observed increased expression and oligomerization of TIR1 into granular structures in muscle of C. elegans due to ROS generated under anaesthesized state. We also observed the formation of lipofuscin granules under anaesthesized condition **(Figure 6 G)**. Lipofuscin granules are autofluorescent, nondegradative granular structures that naturally form in the body of C. elegans. Their accumulation rate increases in response to various stressors like heat shock, high glucose diet, and oxidative stress [38]. Since they emit in the 520-550 nm region, they can act as an easily detectable biomarker for stress in the animal’s body. The increased accumulation of lipofuscin granules under anaesthesised condition in C. elegans points to the systemic stresses faced by the animal when exposed to the treatment.

**Figure 6:**
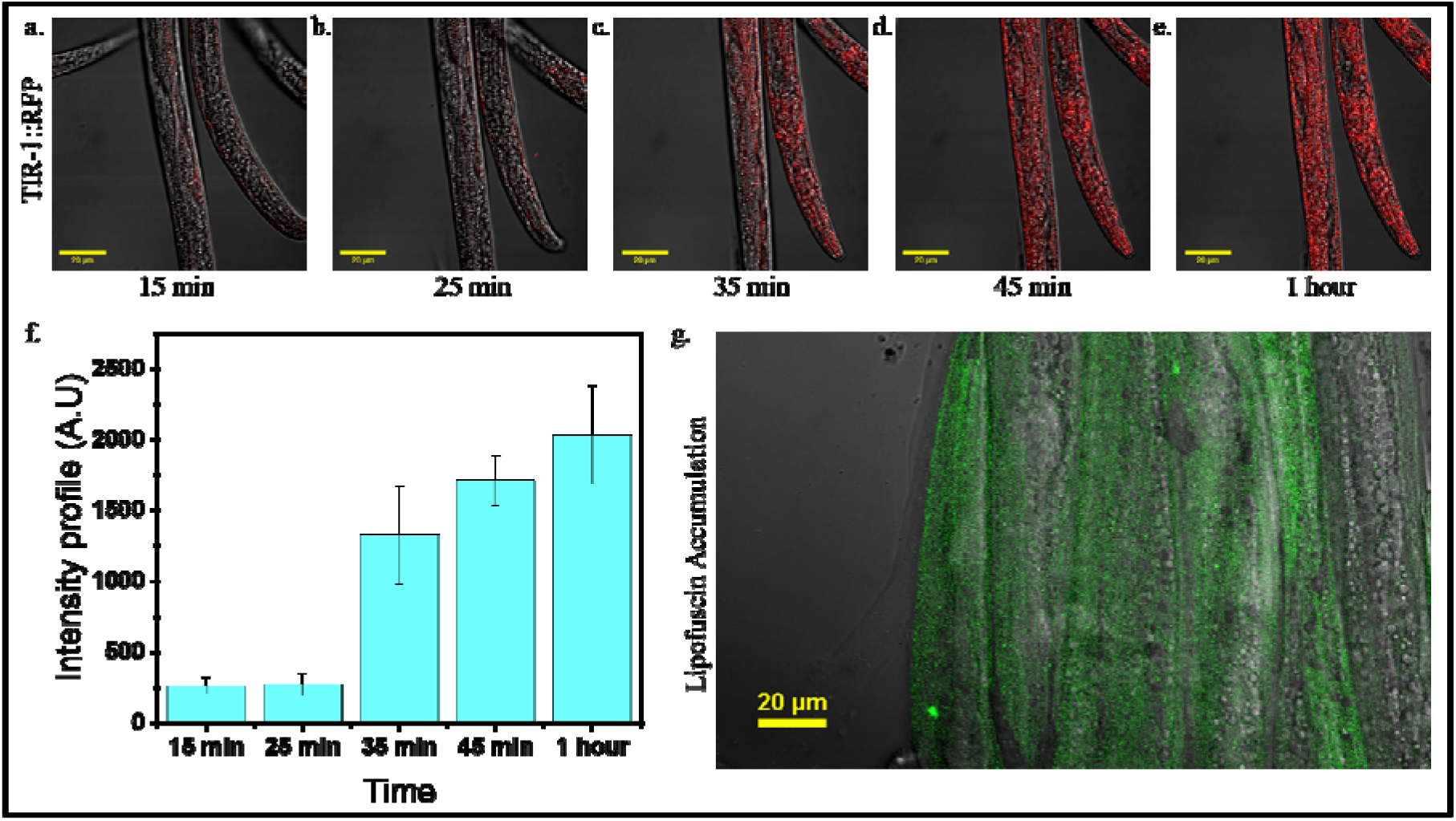
Systemic stress experienced by C. elegans in response to anaesthetic exposure. **(a-e)**. Increase in TIR1::RFP expression and aggregation in muscles of C. elegans ESC360 strain after anaesthetic exposure. **(f)**. Increase in intensity profile of TIR::RFP overtime. **(g)**. Accumulation of lipofuscin granules in the body of C. elegans after long term anaesthetic exposure.

## Discussion

While most anaesthetic research has traditionally focused on neuronal cells due to their clinical relevance in suppressing sensation and consciousness, our study challenges this neurocentric perspective. We demonstrate that anaesthetics such as lidocaine and isoflurane exert widespread and significant effects on non-neuronal cells and subcellular organelles, even at dosages considered clinically safe. These effects are not transient; repeated or prolonged exposure leads to cumulative organellar damage, which may contribute to long-term cellular dysfunction across diverse tissue types. Our findings identify mitochondria, lysosomes, and the nucleus as key vulnerable organelles, significantly expanding the understanding of anaesthetic toxicity beyond the nervous system. In HEK 293T cells, lidocaine exposure resulted in a time-dependent decline in mitochondrial membrane potential, indicative of early mitochondrial dysfunction. This was accompanied by the accumulation of lysosomal aggregates, suggesting an autophagic response likely triggered by rising reactive oxygen species (ROS) levels **(Figure 1)**. Anaesthetics have been shown to inhibit mitochondrial Complex I, impairing the electron transport chain and reducing ATP production [25, 41]. The resultant ROS can damage mitochondrial membranes and mtDNA [41, 42], initiating a cascade of dysfunction that impacts other organelles.

To assess these effects in vivo, we used C. elegans and exposed them to 8% v/v isoflurane, a dosage previously reported as safe [39]. Within 30 minutes, we observed systemic mitochondrial fragmentation and network collapse across tissues **(Figure 2)**, indicating a bioenergetic crisis. This fragmentation compromises ATP generation, disrupting vital processes like transcription, vesicular transport, and survival [40]. Additionally, tubular lysosomes rapidly disappeared following anaesthetic exposure **(Figure 3)**. This loss is likely due to ROS-induced lysosomal membrane permeabilization (LMP), a mechanism previously associated with oxidative stress [43]. The rupture of lysosomes may release hydrolytic enzymes into the cytoplasm, further aggravating cellular damage. At the transcriptional level, anaesthesia did not globally suppress transcription but caused selective downregulation. While RNA Polymerase I-driven transcription remained stable, promoter activity from ttx-3p and t24b8.5p declined significantly within 30 minutes **(Figure 4)**. This suggests a hierarchical transcriptional modulation, likely as an energy-conserving strategy under ATP-limiting conditions [44]. Morphological changes in nuclear structure were also observed. Nuclei became circular and swollen post-anaesthesia, resembling responses seen under osmotic or oxidative stress **(Figure 5)** [45]. This transformation likely results from actin cytoskeleton degradation, which compromises nuclear shape maintenance [46]. Furthermore, his-71::GFP (histone H3.3) fluorescence was reduced in intestinal nuclei but not in muscle nuclei, pointing to tissue-specific resilience or sensitivity **(Figure 5)**. Finally, we confirmed systemic ROS elevation through TIR-1::GFP aggregation and lipofuscin accumulation, both markers of oxidative damage **(Figure 6)**. These findings highlight that anaesthetic exposure activates sustained oxidative stress pathways, which could cause irreversible cellular damage upon chronic or repeated exposure. So, we can say that our study reveals a broad, organelle-centered toxicity profile of anaesthetics, underscoring the need to reevaluate their systemic impact and safety, particularly in non-neuronal and developing tissues.

## Conclusion

Our research presents significant evidence that the anaesthesia toxicity extends beyond neurons, significantly affecting various cell types and different organelles, such as mitochondria, lysosomes, and the nucleus. The effects are systemic, cumulative, and energetically expensive, prompting concerns regarding the long-term cellular consequences in patients undergoing chronic or recurrent anaesthesia, particularly in pediatric or ICU populations. By understanding these processes, we emphasize the necessity for organellar-specific anaesthetic treatments and present novel opportunities for biomarker development to assess anaesthetic-induced cellular stress. Future research may investigate protective medicines or pretreatment techniques to mitigate these systemic cytotoxic effects.

## Materials and Methods

### 1. C. elegans Strain Maintenance

All *Caenorhabditis elegans* strains used in this study were acquired from the Caenorhabditis Genetics Center (CGC), University of Minnesota, supported by the National Institutes of Health. The GL390 strain was used for the visualization of mitochondrial dynamics. The XW8056 strain, which expresses *scav-3::GFP*, was used to observe lysosomal dynamics. The ESC360 strain was employed to monitor cellular stress responses in muscles and RNA polymerase activity. Additionally, the AU78 strain was used to assess the transcriptional efficiency of *TTX-3* and *T24B8*.*5* promoters under anaesthetized conditions.

### 2. Preparation of NGM Plates

All *C. elegans* strains were cultured on nematode growth medium (NGM) agar plates. To prepare NGM, 3 g of NaCl, 2.5 g of peptone, and 17 g of agar were mixed in 975 ml of distilled water and autoclaved. After cooling to approximately 50□°C, 25 ml of 1 M KPO_4_ buffer (prepared using 108.3 g KH_2_PO_4_ and 35.6 g K_2_HPO_4_ in 1 L distilled water), 1 ml of 1 M MgSO_4_, and 1 ml of 1 M CaCl_2_ were added. Before plating, 1 ml of 5 mg/ml cholesterol (dissolved in ethanol) was added when the media temperature was between 37–45□°C to prevent thermal degradation. A total of 10 ml of the final media was poured into each 60 mm Petri plate (Tarsons, 460062).

### 3. Preparation of Bacterial Food Source

*Escherichia coli* OP50 was used as the standard food source for feeding *C. elegans*. The bacteria were grown in Luria Broth (LB) medium prepared by dissolving 20 g of LB powder in 1 L of distilled water, followed by autoclaving and cooling. The medium was inoculated with OP50 and incubated overnight at 37□°C in a shaking incubator. Subsequently, 300 µl of the overnight culture was spread evenly on prepared NGM plates and dried prior to use.

### 4. Inoculation of C. elegans Strains

Adult *C. elegans* were transferred by washing a small area of the NGM plate using M9 buffer. The M9 buffer was prepared by combining 3 g KH_2_PO_4_, 7.52 g Na_2_HPO_4_, 5 g NaCl, and 1 ml of 1 M MgSO_4_ in 1 L of distilled water. The suspended worms were collected and transferred to freshly prepared NGM plates seeded with OP50 bacteria.

### 5. Age Synchronization

For synchronization, Day 3 adult *C. elegans* were washed off the plates using 2 ml of M9 buffer, transferred to centrifuge tubes, and spun at 2000 RPM for 2 minutes in an Eppendorf 5810 R centrifuge. The supernatant was discarded, and the pellet was resuspended in a bleaching solution containing 1 ml distilled water, 500 µl of 1 M NaOH, and 400 µl of NaOCl. The mixture was vortexed for 5 to 8 minutes until the adult worm bodies disintegrated, releasing the embryos. The embryos were collected by centrifugation at 2000 RPM for 2 minutes, washed twice with M9 buffer, and plated onto fresh NGM plates seeded with OP50.

### 6. Anaesthesia Application in C. elegans

Isoflurane was used at 8% v/v concentration to anaesthetize *C. elegans*. The worms were placed in a drop of M9 buffer on a live imaging plate. The drop was surrounded by a thin film of 1% agar to prevent drying during imaging. Isoflurane was introduced through a small perforation in the plate lid, and the setup was immediately sealed using multiple layers of parafilm. Worms exhibited frantic movement initially, followed by complete immobilization within 10 minutes of anaesthetic exposure.

### 7. Cell Maintenance

HEK 293T cells used in the experiments were obtained from NCCS Pune. Cells were cultured at 37□°C and 5% CO_2_ in a Haier HCP-168 incubator. The passage number used for experiments was 16. Standard cell culture protocols were followed for maintenance and sub-culturing.

### 8. Cell Imaging and Anaesthesia

For live-cell imaging, HEK 293T cells were incubated with LysoTracker Red or MitoTracker Green at a concentration of 100 nM for 1 hour. After incubation, the cells were washed and replenished with fresh media before imaging. Lidocaine was used as a liquid anaesthetic at a concentration of 0.02% v/v. Due to its liquid nature, no special sealing of the imaging chamber was necessary during anaesthesia application.

### 9. Confocal Microscopy

Confocal imaging was performed using a Nikon Eclipse Ti inverted microscope, and image acquisition was conducted using Nikon NIS-Elements software. A 488 nm excitation laser with an emission filter range of 525–550 nm was used to visualize GFP and MitoTracker signals, while a 561 nm excitation laser with an emission filter range of 625–650 nm was used to visualize RFP and LysoTracker signals.

## Acknowledgements

We are also grateful to the Indian Knowledge System and Mental Health Applications (IKSMHA) Centre for providing both funding and instrumentation support. We acknowledge the Ministry of Education (MoE) for providing the research stipend.

## Notes

### Competing Interest Statement

The authors have declared no competing interest.

